# The impact of speaker accent on discourse processing: a frequency investigation

**DOI:** 10.1101/2023.12.19.571836

**Authors:** Trisha Thomas, Clara D. Martin, Sendy Caffarra

## Abstract

Previous studies show that there are differences in native and foreign speech processing (Lev-Ari, 2018) while mixed evidence has been found regarding differences between dialectal and foreign accent processing (see: Adank et al., 2009; Floccia et al. 2006 but see also: Floccia et al., 2009; Girard et al., 2008). Within this field, two theories have been proposed. The Perceptual Distance Hypothesis states that the mechanisms underlying dialectal accent processing are attenuated versions of those of foreign (Clarke & Garrett, 2004). While, the Different Processes Hypothesis argues that the mechanisms of foreign and dialectal accent processing are qualitatively different (Floccia et al, 2009). A recent study looking at single-word EEG data, suggested that there may be flexibility in processing mechanisms (Thomas et al., 2022). The present study deepens this investigation by addressing in which frequency bands native, dialectal and foreign accent processing differ when listening to extended speech. Electroencephalographic data was recorded from 30 participants who listened to dialogues of approximately six minutes spoken in native, dialectal and foreign accents. Power spectral density estimation (1-35 hz) was performed. Linear mixed models were done in frequency windows of particular relevance to discourse processing. Frequency bands associated with phoneme [gamma], syllable [theta], and prosody [delta] were considered along with those of general cognitive mechanisms [alpha and beta]. Results show power differences in the Gamma frequency range. While in higher frequency ranges foreign accent processing is differentiated from power amplitudes of native and dialectal accent processing, in low frequencies we do not see any accent-related power amplitude modulations. This suggests that there may be a difference in phoneme processing for native accent types and foreign accent, while we speculate that top-down mechanisms during discourse processing may mitigate the effects observed with short units of speech.

## 1. Introduction

In our current society of global mobility, the study of speech accent is more relevant than ever. Many previous studies have shown impaired speech comprehension of unfamiliar or accented speech due to the unique challenges that non-native speech presents the listener (Floccia et al., 2006; Munro and Derwing, 1995; Schmid and Yeni-Komshian, 1999; Anderson-Hsieh and Koehler, 1988; Major et al., 2002). As cross-cultural interactions become increasingly common, it is important to investigate the underlying mechanisms involved in processing non-standard speech in its multiple forms. Most studies focus on contrasting foreign accent from native accent using behavioral or evoked responses from electrophysiological (EEG) methods. Here we use a less common EEG technique to study how our brain responds to not only native and foreign accent but also dialectal accent.

While foreign accent is often the focus when discussing non-standard speech, dialectal speech can also be considered non-standard speech from the perspective of a native listener. A dialectal accent would come from a native speaker of the same language of the listener who is from a different county or geographical region. These accent types possess native phoneme variations, distinguishing them from foreign accents that often contain non-native variations resulting from the sounds or syntactic rules of the native language of the speaker. Because of these differences, this study aims to understand how dialectal accent is processed in relation to native and foreign accent.

It is generally agreed upon that foreign accent presents the greatest challenge to the listener as compared to native accent. Foreign accents are often identified more readily than dialectal accents and have a greater impairment on lexical retrieval (Girard et al., 2008; Floccia et al., 2009). However, the case of dialectal accent is still understudied and it is not clear what is its impact on brain responses relative to the foreign and the native accent. The findings of Girard and colleagues (Girard et al., 2008) and Floccia and colleagues (Floccia et al., 2009) seem to be in line with the Different Processes Hypothesis, which suggests that there are qualitative differences in the processing mechanisms recruited for dialectal and foreign speech normalization (Floccia et al., 2009). In this case, the main qualitative distinction is between accents that share a native language and non-native accents. Another hypothesis, the Perceptual Distance Hypothesis, suggests that accents can be placed on a perceptual scale based on their acoustic distance from native speech and thus dialectal accent is processed more like an attenuated version of foreign accent (Clarke & Garrett, 2004). Some behavioral studies have provided support for this hypothesis, with Adank and colleagues (Adank et al., 2009) reporting decreased processing speed as accents become less familiar or more foreign. An earlier study by Floccia and colleagues (Floccia et al., 2006) also provided evidence for this hypothesis when they found slower word recognition for words uttered in an unfamiliar dialectal accent, and even more slowly in a foreign accent.

In addition to behavioral techniques, EEG methods have been employed to investigate accented speech comprehension due to their ability to provide valuable time-sensitive information. One common technique for investigating the effects of accent on speech processing using electrophysiological methods is the Event-Related Potential (ERP) analysis. Event-related potentials (ERPs) are a type of neurophysiological measurement technique created by signal averaging used to study brain activity in response to specific events or stimuli. ERPs related to speech processing have shown that the time course of speech analysis changes as a function of accent.

Early evoked responses typically associated with phonological analysis have mainly shown differences between foreign and native accent, supporting the Different Processes Hypothesis (see Thomas et al., 2022; Goslin et al., 2012). In late evoked responses the results are more mixed, with some studies supporting the Perceptual Distance Hypothesis (see Jiang et al., 2020) and others the Different Processes Hypothesis (see Hanulíková et al., 2012). While ERP studies have been instrumental in understanding differences between processing mechanisms of native and non-native accented speech, because this technique reduces the complexity of the electrophysiological signal to one time-dependent variable, it does not allow for tracking multiple rhythmic processes occurring in parallel. Therefore, it is also useful to focus on the frequency domain of the EEG signal when studying linguistic processes, especially in the case of sentences and discourses where the rhythmic activity of the brain can be tracked over large time windows.

While time-based approaches, such as ERP analysis, can provide us with beneficial information, time-frequency methods offer two main advantages. They provide information about electrophysiological activity in a naturalistic situation (passive listening to continuous speech) and they are able to detect electrophysiological brain activity undetectable by ERPs, reflecting different parallel oscillatory patterns that are not time-locked to the presentation of a stimulus. Oscillations are thought to reflect the coordinated activity of large groups of neurons and may play a critical role in synchronizing neural processing across different brain regions. These oscillatory patterns are generally categorized according to their frequency bands: delta (δ < 4 Hz), theta (θ 4-8 Hz), alpha (9-12 Hz), beta (13-25 Hz) and gamma (> 25 Hz). Power spectral density estimation is a time-frequency analysis that calculates the distribution of power in different frequency bands. Previous studies have linked different frequency bands to different aspects of speech processing, such as phonetic analysis, semantic processing, and syntactic integration.

In the field of linguistic research, three frequency bands have been characteristically linked to linguistic properties of speech: the delta, theta and gamma bands (for review, see (Giraud & Poeppel, 2012); (Grabot et al., 2017); (Meyer et al., 2017)). To a lesser extent, some research also has correlated alpha and beta band power with linguistic processes.

Within low-frequency ranges (i.e., δ and θ), it has been long suggested that theta oscillations reflect syllable tracking. Peña and Melloni found that theta power was higher when listening to forward utterances as opposed to backward ones (which disrupt the syllabic structures of the utterance), suggesting that theta power is involved in tracking syllable patterns ((Peña & Melloni, 2012)). Several studies have provided evidence that theta oscillations synchronize with syllable onset ((Luo & Poeppel, 2007); (Howard & Poeppel, 2012); (Peelle et al., 2013); (Doelling et al., 2014)) and thus aid in the identification of syllable boundaries. Limited additional studies have also suggested that theta oscillations may be additionally involved in lexical-semantic retrieval ((Bastiaansen et al., 2008) and syntactic processing ((Bastiaansen et al., 2002)). Delta bands have been linked to intonation or prosodic processing due to their phase coherence with the pitch contour of speech (Giraud & Poeppel, 2012); (Bourguignon et al., 2013); (Mai et al., 2016)).

Within high-frequency ranges (i.e., β and γ), more previous research has been done on gamma oscillations’ role in speech processing. Traditionally, synchronization of the gamma band amplitude has been suggested to reflect phonemic-categorical perception ((Lehongre et al., 2011)). Ortiz-Mentilla and colleagues found that even as early as 6 months of age, high gamma power is enhanced during the discrimination of native phoneme contrasts, suggesting that it plays an early role in the perception and categorization of phonemic features, allowing for preferential processing of native phoneme contrasts ((Ortiz-Mantilla et al., 2013)). While low gamma synchronization has been linked to acoustic processing ((Gross et al., 2013)). The Beta frequency band has also been correlated with linguistic processes. It has been shown to decrease in situations of mismatch between semantic predictions and reality ((Lewis & Bastiaansen, 2015); (Lewis et al., 2016) see also Weiss & Mueller, 2003 for beta band power role in semantics)). The mid-range alpha frequency, however, is mostly attributed to more general cognitive functions and has been classically related to the inhibition of task-irrelevant information (for review, see (Klimesch et al., 2007)) and suppression of cortical excitability (Jensen and Mazaheri 2010). Despite alpha being associated most prominently with generalized top-down mechanisms, some previous works have provided evidence that it may play a role relevant to sentence comprehension in verbal working memory (Krause et al., 1996); (Maltseva et al., 2000); (Jensen et al., 2002); (Sauseng et al., 2005); (Leiberg et al., 2006)).

While there are many studies linking various neural oscillations to linguistics processes, very little work has been done to differentiate dialectal and foreign accents in the frequency domain. While behavioral and ERP studies have provided evidence for both previously proposed hypotheses, this is the first study to our knowledge comparing neural oscillations of foreign, dialectal and native accent perception. Examining discourse through the frequency domain allows us to see how speech accent affects brain rhythmic activities at different frequency ranges thus enabling us to understand the robustness of previously observed evidence in favor of either the Perceptual Distance or Different Processes Hypothesis.

### 1.1. The Present Study

The present study aims to investigate how the brain processes accented speech across different frequency bands using electrophysiological methods. We critically evaluate the aforementioned previously proposed hypotheses of accented speech processing in order to clarify the processing of foreign, dialectal and native accented speech. Special attention is placed on disentangling how we process dialectal accent from both native and foreign accent during the listening of short stories. Analyses include Power spectral density estimation for each accent condition because of its relevance to EEG analysis of extended speech. Specifically, we focus on Delta, (1-3 Hz) Theta (4-8 Hz) and Low Gamma (25-35 Hz) waves, associated with prosodic, syllabic and phonemic processing, respectively (see Meyer, L., 2017 for a review).

According to the Perceptual Distance Hypothesis, we should observe a gradient in the average power amplitude of frequency bands that are implicated in the processing of native, dialectal and foreign accent, respectively. Conversely, results showing that dialectal and native accents are processed similarly would support the Different Processes Hypothesis. Thus, we would see a categorical discrimination between the foreign accent and the other more standard accents (native and dialectal).

## 2. Materials & Methods

### 2.1. Participants

Thirty Spanish natives^1^ participated in the study. One participant was excluded due to low performance on the target detection task (see method section for further explanation of this task; mean accuracy of this participant: 37.5%). Another participant was excluded for low comprehension question performance (mean accuracy: 53.3%). The final sample of participants consisted of 28 females^2^ (mean age: 22.7 years, SD: 3.56, age range: 19-31 years, Spanish age of acquisition: 0). All participants lived in the Basque country and considered the Basque-Spanish accent as their native accent. All participants were right-handed and had normal or corrected-to-normal hearing and vision. No participant reported a history of neurological disorders. All participants signed an informed consent form before taking part in the study that was approved by the Basque Center on Cognition, Brain and Language ethics committee. They received monetary compensation for their participation.

### 2.2. Materials

Three dialogues were recorded by six female speakers in their thirties with differing accents. Each dialogue was recorded by two speakers of the same accent (native, foreign, dialectal). Each pair of same-accent speakers recorded the three stories (total of recorded stories: 9). Story type and pair of speakers were fully counterbalanced so that each participant listened to dialogues produced by each pair of speakers without repetition of the same story (total of 3 stories presented). Stories were recorded in a Basque-Spanish native accent, a Cuban-Spanish accent and an Italian-Spanish accent (i.e., hereafter native, dialectal and foreign accent, respectively). The native speakers were born and lived in Spain (Basque country). The dialectal and foreign speakers were chosen for their strong accents and high level of Spanish (in the case of the Italian speakers). Overall, the recordings did not significantly differ in duration (ms) across accents [foreign: 519.7, SD:128.7; native: 505.7, SD: 137.2; dialectal: 519.9, SD:131.6; one way ANOVA: (F(2,117)=0.15, p =0.86)]. Accent strength ratings were collected from a separate normative study consisting of eleven participants (average age =23, SD=10) who completed a short online survey where they listened to clips of each accent and rated the accent strength from 1 (mild accent) to 5 (strong accent). Results showed a clear effect of accent (one way ANOVA: F(2,10)=6.01, p=0.009). Follow-up analyses of the clip ratings corrected with the Fisher’s Least Significant Difference (LSD) showed that the dialectal accent was marginally significantly different from the native accent (t(10)=2.13, p=.06), the foreign accent was significantly different from the native accent (t(10)=3.16,p<.001) and the dialectal and foreign accents were not rated significantly different from each other in terms of strength (t(10)=1.15, p=.28).Ten comprehension questions for each dialogue were also created^3^.

### 2.3. Procedure

Participants carried out a blocked-design experiment consisting of 4 blocks, three speech blocks and one silence block^4^. Participants were seated in a sound-attenuated room in front of a computer screen and were asked to listen to narrative dialogues (or silence) and occasionally perform a target detection task to maintain their attention by pressing the spacebar with both index fingers when they heard certain target words. After each block, they filled out a questionnaire with 10 comprehension questions about the dialogue. Each block consisted of a dialogue (or silence) presented through speakers while a fixation cross was displayed on the screen throughout the entirety of the dialogue. The experimental session lasted about an hour. During the silence block, participants still occasionally heard the target words and had to press the spacebar. The block order was counterbalanced across participants.

### 2.4. EEG Data Recording and Time-Frequency Analyses

The EEG signal was recorded from 27 channels placed in an elastic cap: Fp1, Fp2, F7, F8, F3, F4, FC5, FC6, FC1, FC2, T7, T8, C3, C4, CP5, CP6, CP1, CP2, P3, P4, P7, P8, O1, O2, Fz, Cz, Pz (see fig.1). Two external electrodes were placed on the mastoids, two were on the ocular canthi, one above and one below the right eye. All sites were referenced online to the left mastoid. Data were recorded and amplified at a sampling rate of 250 Hz. Impedance was kept below 10 KΩ for the external channels and below 5 KΩ for the electrodes on the scalp. A low-pass filter of 30 Hz and a high-pass filter of 0.01 Hz were applied. Vertical and horizontal eye movements were corrected by performing an Independent Components Analysis (ICA). The fastICA method was used. Time-frequency analysis of continuous EEG data was done with a Morlet wavelet decomposition using MNE software (Gramfort et al., 2013). This method was used to decompose trial time-frequency values between 1 and 35 Hz for the 28 electrodes placed on the scalp (steps = 13). The average total power values were baseline corrected with a log-ratio. We used the pre-target 500 ms silent period (-500 to 0 ms) as baseline. For each accent condition and frequency range, the resulting power was averaged across time and channels.

**Fig 1.**
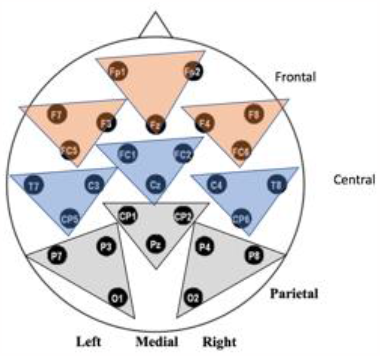
Schematic of electrode montage with topographic organization labeled.

## 3. Results

### 3.1. Behavioral Results

Participants showed average accuracy during the online target detection task meant to test attention (mean overall accuracy: 71.4%, SD:13.2)^5^. Participants showed high accuracy on comprehension question performance (mean overall accuracy: 83.6%, SD:8.7). Importantly, accuracy in target detection (F(2,81)=1.67, p=0.19) and comprehension questionnaire (F(2,82)=0.15, p=0.86) did not significantly differ across the three accent conditions (all p values above 0.05).

### 3.2. Time-Frequency Results

Average power amplitude modulation in the EEG of participants was analyzed with a mixed linear model with participants as random intercept and accent as fixed factor (native, dialectal and foreign). An accent effect was observed only within the gamma frequency range, see figure XX.

**Figure xx.**
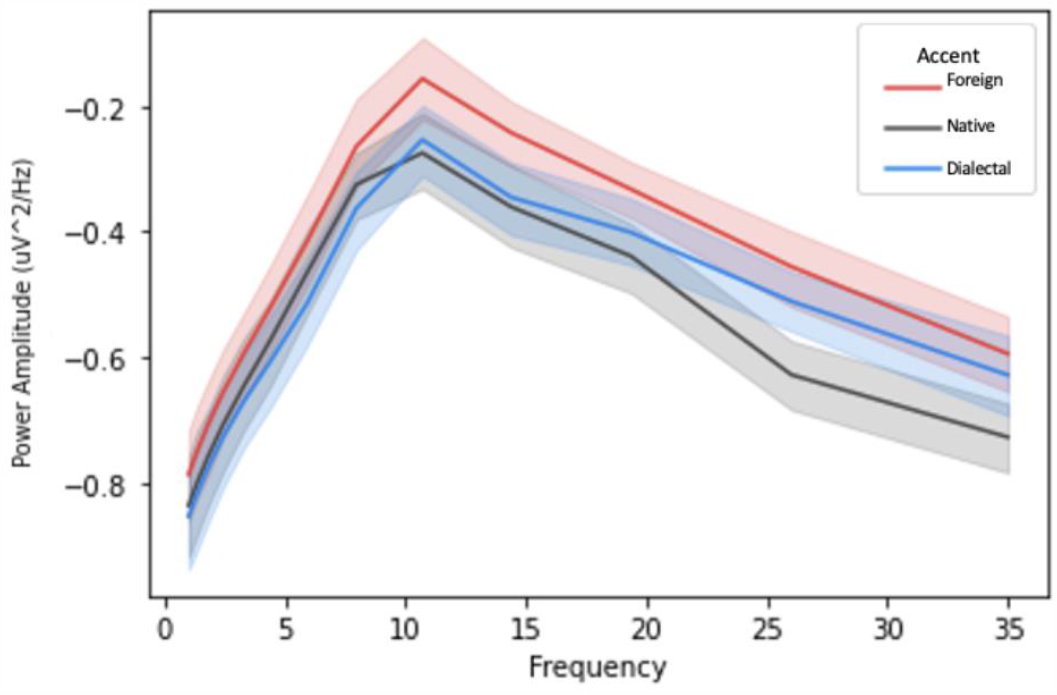
Power amplitude modulation in the EEG across accents.

#### Gamma (25 to 35 hz) phoneme

In gamma frequencies we found a significant difference between native and foreign accents (native vs. foreign: *β*=0.15, *SE*=0.08, *t*=1.97, *p*=0.048). However, no difference was found between dialectal accent and either native or foreign accent (native vs. dialectal: *β*=0.11, *SE*=0.08, *t*=1.40, *p*=0.16; foreign vs. dialectal: *β*=-0.04 *SE*=0.08, *t*=0.58, *p*=0.56; see Figure xx).

**Fig. xx.**
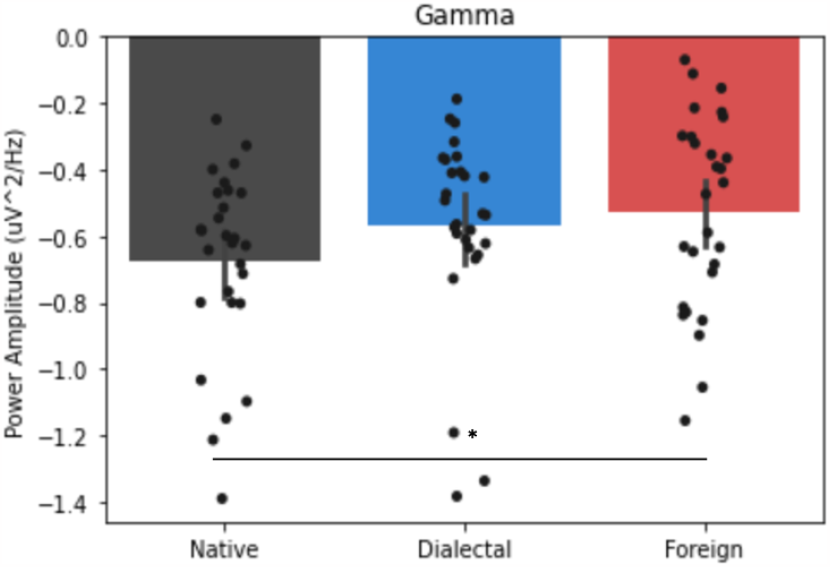
Average Gamma power between accents, ci=95.

In Delta, Theta, Alpha and Beta frequencies, we did not observe any power difference between the three accents (see Table xx, Fig xx).

**Fig. xx.**
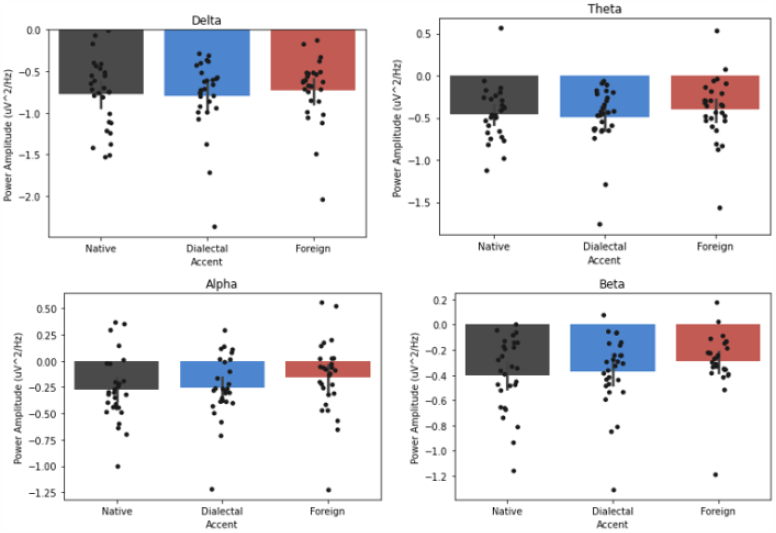
Average Beta power between accents, ci=95.

**Table xx.** Summary of mixed linear model results for each frequency range which show a lack of significant differences between accents.

| Frequency band | Accent contrasts | $\beta$ | $SE$ | $t$ | $p$ |
| --- | --- | --- | --- | --- | --- |
| Delta (1 to 3 Hz)<br>prosody | native vs. dialectal | -0.02 | 0.09 | 0.19 | 0.84 |
|  | native vs. foreign | 0.05 | 0.09 | 0.53 | 0.60 |
|  | foreign vs. dialectal | -0.07 | 0.09 | 0.72 | 0.47 |
| Theta (4 to 8 Hz)<br>syllable | native vs. dialectal | -0.04 | 0.08 | 0.66 | 0.62 |
|  | native vs. foreign | 0.05 | 0.08 | 0.49 | 0.51 |
|  | foreign vs. dialectal | -0.09 | 0.08 | 1.15 | 0.25 |
| Alpha (9 to 12 Hz) | native vs. dialectal | 0.02 | 0.08 | 0.26 | 0.79 |
|  | native vs. foreign | 0.12 | 0.08 | 1.48 | 0.14 |
|  | foreign vs. dialectal | -0.10 | 0.08 | 1.21 | 0.22 |
| Beta (13 to 24 Hz) | native vs. dialectal | 0.03 | 0.07 | 0.39 | 0.69 |
|  | native vs. foreign | 0.11 | 0.07 | 1.66 | 0.10 |
|  | foreign vs. dialectal | -0.08 | 0.07 | 1.27 | 0.21 |

## 4. Discussion

This study aimed to advance previous investigations of online accented speech processing through the lens of oscillatory activity power amplitude fluctuations during listening to different speech accents. We further aimed to better understand the similarities and/or differences between mechanisms of dialectal and foreign-accented speech processing by examining the results in the context of two hypotheses of accented speech processing: the Different Processes Hypothesis and the Perceptual Distance Hypothesis.

Previous studies have mostly been conducted at the level of the single word and have emphasized processing differences between native and foreign speech processing ((Lane, 1963); (Lev-Ari & Keysar, 2012); Lev-Ari,(“Credibility of Native and Non-Native Speakers of English Revisited: Do Non-Native Listeners Feel the Same?,” 2017); (Munro & Derwing, 1995); (van Wijngaarden, 2001)). But whether native listeners process native category variations, i.e. dialectal accent, according to acoustic distance from native speech (Perceptual Distance Hypothesis) or with ‘nativeness’ as a processing category (Different Processes Hypothesis) is less clear.

This study examined these hypotheses by investigating how the brain processes accented speech across different frequency bands using electrophysiological methods. We focused on Delta, (1-3 Hz) Theta (4-8 Hz) and Low Gamma (25-35 Hz) waves, because of their previous association with prosodic, syllabic and phonemic processing, respectively. We also considered Alpha and Beta because of their relation to attentional processing and top-down mechanisms.

We found a significant difference between the power amplitude of native and foreign accent in the low gamma frequency range with no clear evidence on whether dialectal accent is differentiated from native accent. Because of this, we ran an exploratory follow-up Bayesian analysis to understand the natureal of the null effect between dialectal and native accent. This analysis provided weak support for the null hypothesis (BF: 0.89) and thus evidence that dialectal accent phoneme processing is similar to native accent phoneme processing.This finding shows that unique processing mechanisms are required for the phonological processing of foreign accent throughout discourse processing as compared to native accent types. This provides evidence for the Different Processes Hypothesis by supporting differential processing for non-native accent but not for dialectal accent as compared to native accent. This is perhaps due to the ‘nativeness’ of the deviations in dialectal pronunciation, meaning that ‘coherent deviations’ (see (Wells, 1982); (Goslin et al., 2012)) of the speech in a dialectal accent are easily adapted to by listeners while foreign deviations make acoustic extraction more difficult.

These findings seem to somewhat align with previous ERP studies of accent processing, including a previous study on single word processing (see (Thomas et al., 2022) but see (Goslin et al., 2012)). In Thomas et al.’s study, we found a difference between foreign and both native accent variations (native and dialectal) at early ERP components associated with the extraction of acoustic features. Similarly, at ERP components associated with lexico-semantic integration, we no longer observed accent differences.

The results of the present study similarly suggest that non-native accent affects early stages of phoneme processing both at single word and discourse levels. They further suggest that these effects are uniquely seen for non-native accents rather than unfamiliar native variations.

These results contribute to the literature supporting the Different Processes hypothesis and further support the idea of a binary native/non-native processing mechanism in the more naturalistic setting of extended discourse. Furthermore, accent seems to have the largest effect on phonological analysis while successfully resolved in the semantic processing phase.

Previous studies show that listeners employ top-down resources to process speech in challenging situations, engaging contextual cues, predictive processing, and prior knowledge about the topic to enhance comprehension ((Dave et al., 2021); (Foucart et al., 2015); cf. (Schiller et al., 2020)). However, in the alpha frequency band, associated with top-down mechanisms and attentional control/inhibition, we did not observe any significant effects. This is surprising in light of the previous findings, as we may expect to see some engagement of inhibitory processing in attentionally demanding situations, especially given the lingering effects of both dialectal and foreign accent-delivered speech on memory. Thus, whether accent is truly resolved at the phonological level or simply mitigated by the engagement of top-down strategies remains to be further explored and should be additionally scrutinized in future studies.

### 4.1. Conclusion

We found that while in higher frequency ranges, power amplitudes of foreign accent processing are differentiated from power amplitudes of native accent processing, in low frequencies we do not see any accent-related power amplitude modulations. This suggests that while the native/non-native accent distinction modulates the way we process speech at the phonological level, it does not seem to have such an effect at higher levels of processing (i.e., lexicon and semantics).

## Acknowledgments

This research was supported by the Basque Government through the BERC 2022-2025 program and by the Spanish State Research Agency through BCBL Severo Ochoa excellence accreditation CEX2020-001010-S. The research was also supported by the Spanish Ministry of Economy and Competitiveness (PID2020-113926GB-I00 to C.D.M.), and the European Research Council (ERC) under the European Union’s Horizon 2020 research and innovation programme (grant agreement No 819093 to C.D.M.; No 837228 to SC). This work was also supported by the Basque Government [PIBA18-29 to CDM]; the Italian Ministry of University and Research (Programma giovani ricercatori Rita Levi Montalcini; (R)FARORIENTED22; P2022SMEJW) to SC; and the Programa Predoctoral de Formación de Personal Investigador No Doctor del Departamento de Educación del Gobierno Vasco (PRE_2021_2_0006 to TT).

## Data availability statement

Code and data to reproduce the reported findings are available at [INSERT URL UPON PUBLICATION]

## Author contributions

All authors contributed to the conceptualization of the study. SC and CDM performed the data collection. TT performed the data analysis and carried out the initial manuscript preparation. TT, CDM, and SC collaborated on the final version of the manuscript.

## Ethics approval statement

The study was approved by the Basque Center on Cognition, Brain and Language Ethics Committee.

## Appendix List of comprehension questions Dialogue 1

(English)

1. During what time of day does the dialogue take place?
2. What is Emma trying to open?
3. What jobs do the characters have?
4. Who does Emma want to seem like?
5. What temperature is it outside?
6. Who had warned Sara that she should distrust a certain person?
7. What is the word that begins with R and that scares Sara?
8. While they are talking, is the lamp on?
9. What do the ladies of the house embroider?
10. What do the characters eat for breakfast?

Dialogue 2 (English)

1. Where does Anna want to go?
2. What type of transportation does Anna say she will go on?
3. Where can single women in the city work?
4. What do they give to women who work in the city?
5. In what century does the conversation take place?
6. Who is Anna in love with?
7. Does Anna have parents?
8. What would Anna buy with the money saved?
9. Who will read the letter to Elena?
10. What would happen to Elena if something bad happened to Anna?

Dialogue 3 (English)

1. What was Carla afraid of?
2. What was the woman from the house across the street doing?
3. What does Carla offer María to drink?
4. What is the name of the boy who was in the house before?
5. Did Mary’s brother go to Israel 10 years ago?
6. What is the name of María’s brother?
7. What would warm your body and bring you joy?
8. Where does Carla work?
9. How does María feel after chatting with Carla?
10. What characters do Carla and María invent while having tea?

## Footnotes

1 Due to their geographical location in the Basque Country, all participants were also fluent in Basque as early L2

2 Only females were recruited for this study in order to avoid cross-gender listening effects (Banaji and Hardin, 1996; Cacciari and Padovani, 2007)

3 See appendix for list of questions

4 This was added as a potential baseline in case the EEG of our preferred baseline (the pre-target silent segment) would have been too noisy

5 Accuracy was not very high, but note that participants had to pay attention to the information provided in the dialogue together with tracking the words amigo(s)/amiga(s) which accounts for an accuracy not approaching ceiling.

